# Diatom ultrastructural diversity across controlled and natural environments

**DOI:** 10.1101/2025.06.16.659906

**Authors:** Serena Flori, Felix Mikus, Eliott Flaum, Kevin Moog, Sarah Guessoum, Thomas Beavis, Soraya M. Zwahlen, Inés Romero-Brey, Viola Oorshot, Marine Olivetta, Melissa Steele-Ogus, Ellen Yeh, Mobile Labs Team, Omaya Dudin, Yannick Schwab, Gautam Dey, Flora Vincent

## Abstract

Diatoms are ubiquitous aquatic microalgae critical to our planet, that were amongst the pioneer model organisms in cell biology for their large and transparent cell structure. However, their robust silica cell wall renders diatoms impermeable to many dyes and antibodies, and complicates the intracellular delivery of gene editing tools - driving in part the eventual decline of diatoms as mainstream model species despite their unique cellular physiology and remarkable ecological success. Here, we demonstrate that cryo-fixation combined with ultrastructural expansion microscopy (cryo-ExM) can overcome the silica barrier across diverse diatom species spanning over 80 million years of evolutionary time. We illustrate the potential of cryo-ExM to provide scalable, cost-effective volumetric imaging of diatom ultrastructure in laboratory cultures, as well as field-collected samples from the pan-European TREC expedition. We first reveal striking similarities in interphase microtubule organization across diverse diatom species by characterizing cytoskeletal arrangements throughout cell cycles and populations, uniting both pennate and centric morphologies under shared principles. We further unveil diatom photosynthetic diversity through qualitative and quantitative comparative analysis of chloroplast and pyrenoid morphologies, demonstrating that each diatom species architects unique photosynthetic machinery. Using cryo-ExM on environmental samples further exposes intricate diatom symbioses, revealing tight spatial organisation of ecological interactions. This methodology makes diatoms more accessible for modern and comparative cell biology research, providing new opportunities to investigate the cellular mechanisms of one of Earth’s most successful photosynthetic groups.

**Highlights:**

- Cryo-ExM successfully overcomes diatom frustule barriers to achieve consistent, high-resolution immunofluorescence across evolutionarily diverse species - allowing for comparative cell biology in one of the most important phytoplankton groups on the planet.
- Characterizing microtubule organisation in asynchronously cycling single cells and colonial chains, reveals core features of interphase microtubule organisation across multiple pennate and centric diatom species.
- Comparative analysis of chloroplast and pyrenoid morphologies across species provides qualitative and quantitative insights on photosynthetic diversity, suggesting that photosynthetic architectures are unique to each diatom.
- Applied to environmental samples, cryo-ExM provides detailed insight on spatial organisation of diatom symbioses.

## Introduction

Diatoms are aquatic photosynthetic microeukaryotes responsible for 20% of annual primary productivity and the cornerstone of marine ecosystems (Field et al., 1998). They supply large amounts of dissolved oxygen to marine life, support the entire food chain as pasture of the sea and export carbon to the bottom of the ocean through aggregation and sinking (Armbrust, 2009). Diatoms are considered one of the most successful groups of eukaryotic microalgae on Earth (Malviya et al., 2016; Pierella Karlusich et al., 2025), with tens of thousands of described species that have colonized both freshwater and marine aquatic environments, sediments, and even soils. Yet, our knowledge of diatom cell biology remains restricted to a handful of cultivated species, despite unique biological features shared by almost all species in the clade, such as their sturdy and characteristic silica cell wall or size-induced sexual reproduction.

Diatoms were amongst the first model organisms used to study cell biology in the late 19^th^ century, because their large, clear cells made them ideal for early light microscopic observations, especially for dynamic processes like cell division, migration and photosynthesis (Cande & McDonald, 1985; Lauterborn, 1896; J. Pickett-Heaps, 1991; J. D. Pickett-Heaps & Tippit, 1978). For instance, early studies revealed the presence of a multi-layered acentriolar microtubule-organising centre (MTOC), but the molecular composition, duplication cycle and precise role of this structure in orchestrating mitotic spindle dynamics remain unknown to date. Immunofluorescence and light microscopy were used to study the role of kinesin motors in *Cylindrotheca fusiformis* cell division (Hogan et al., 1993) as well as the function of cell surface proteoglycans in *Stauroneis decipiens* gliding motility (Lind et al., 1997). Live-cell imaging with endogenous and overexpressed GFP-tagged and YFP-tagged proteins has been used to study silica morphogenesis in *Thalassiosira pseudonana* (Scheffel et al., 2011), gliding in *Craspedostauros australis* (Poulsen et al., 2023) and cell division in *Phaeodactylum tricornutum* (Tanaka et al., 2015). Later on, and to gain higher spatial resolution, antibody labelling was coupled to electron microscopy to study photosynthesis, and localize protein complexes such as photosystems in thylakoid membranes (Flori et al., 2017). More recently, the visualisation of small sections of native, frozen cells at near-atomic resolution using cryo-electron tomography (cryo-ET) has led to the discovery of a protein shell localized in the chloroplast and surrounding the pyrenoid (the “PyShell”), that plays a critical role in carbon concentrating mechanisms (Nam et al., 2024; Shimakawa et al., 2024). Despite these noteworthy successes in probing diatom cell biology with imaging, the field faces a critical limitation: the robust silica cell wall of diatoms presents a significant barrier to the penetration of most antibodies and dyes. Consequently, while diatoms remained valuable for research areas like biomineralization or ecology, their use in mainstream cell biology declined as more tractable systems like yeast, mammalian cells, and *Arabidopsis* came to dominate the field (De Martino et al., 2009).

Here, we unlock the potential of diatom cell biology research with ultrastructural expansion microscopy (U-ExM) across lab- and field-based samples. U-ExM is a super-resolution imaging technique that physically enlarges cells and tissues within a swellable hydrogel to achieve nanoscale resolution using conventional light microscopes (Gambarotto et al., 2019). While U-ExM has worked on a variety of species including cell walled ones, allowing for immunolabeling where previously unfeasible (Mikus et al., 2024; Olivetta et al., 2024; Shah et al., 2024), diatoms have so far resisted attempts to expand. Combining cryo-fixation with U-ExM, protocol named cryo-ExM (Laporte et al., 2022), we establish a working expansion protocol effective across diatom cultured species that diverged 80 million years ago, achieving an expansion factor of roughly 4-fold and targeting proteins essential to the cell cycle and host photosynthetic machinery. By performing cryo-ExM on diatoms collected during the pan-European expedition TREC, we successfully preserve potential symbiotic associations that often underlie diatom interactions with their environment (Vincent & Bowler, 2020, 2022). We expose hidden cytoskeletal unity across diatom diversity by mapping microtubule organization throughout cell cycles and populations, discovering shared interphase principles that bridge pennate and centric morphologies. Our comprehensive morphological analysis then reveals how each diatom species crafts specialized photosynthetic machinery through distinct chloroplast and pyrenoid architectures. Taken together, the application of cryo-ExM to diatoms offers unprecedented opportunities to dive into the cellular underpinnings of one of the most successful groups of photosynthetic organisms on the planet.

## Results and discussion

### Cryo-expansion microscopy enables robust immunofluorescence analysis in diverse diatoms from cultures and environmental samples

The ornamented silica cell wall of diatoms, called the frustule, is at the basis of diatom taxonomy, separating them into centric (circular symmetry) and pennate (bilateral symmetry) species. However, this rigid sturdy material also complicates the applicability of many immunofluorescence techniques that often rely on the penetration of antibodies, including standard, proteinase K based ExM protocols (Chen et al., 2015). Therefore, we decided to use the alternative U-ExM protocol, which allows for immunofluorescence after sample expansion, combined with the cryo-ExM approach that replaces chemical fixation with vitrification via plunge freezing (Laporte et al., 2022). We used NHS ester, which preferentially labels dense proteinaceous structures, as a general ultrastructural marker (M’Saad & Bewersdorf, 2020). We show that labelling and resolution obtained using standard confocal systems improves significantly post-expansion (**Fig. 1A**). When comparing cells before and after expansion we obtained an expansion factor of 4.3-fold for *Coscinodiscus granii* (**Fig. 1B**) and 4.1-fold for *Chaetoceros neogracilis* (**Supp. Fig. 1 A,B**). This is in line with previously reported expansion factors across multiple systems (Hinterndorfer et al., 2022; Langner et al., 2024; Liffner et al., 2024; Mikus et al., 2024).

**Figure 1:**
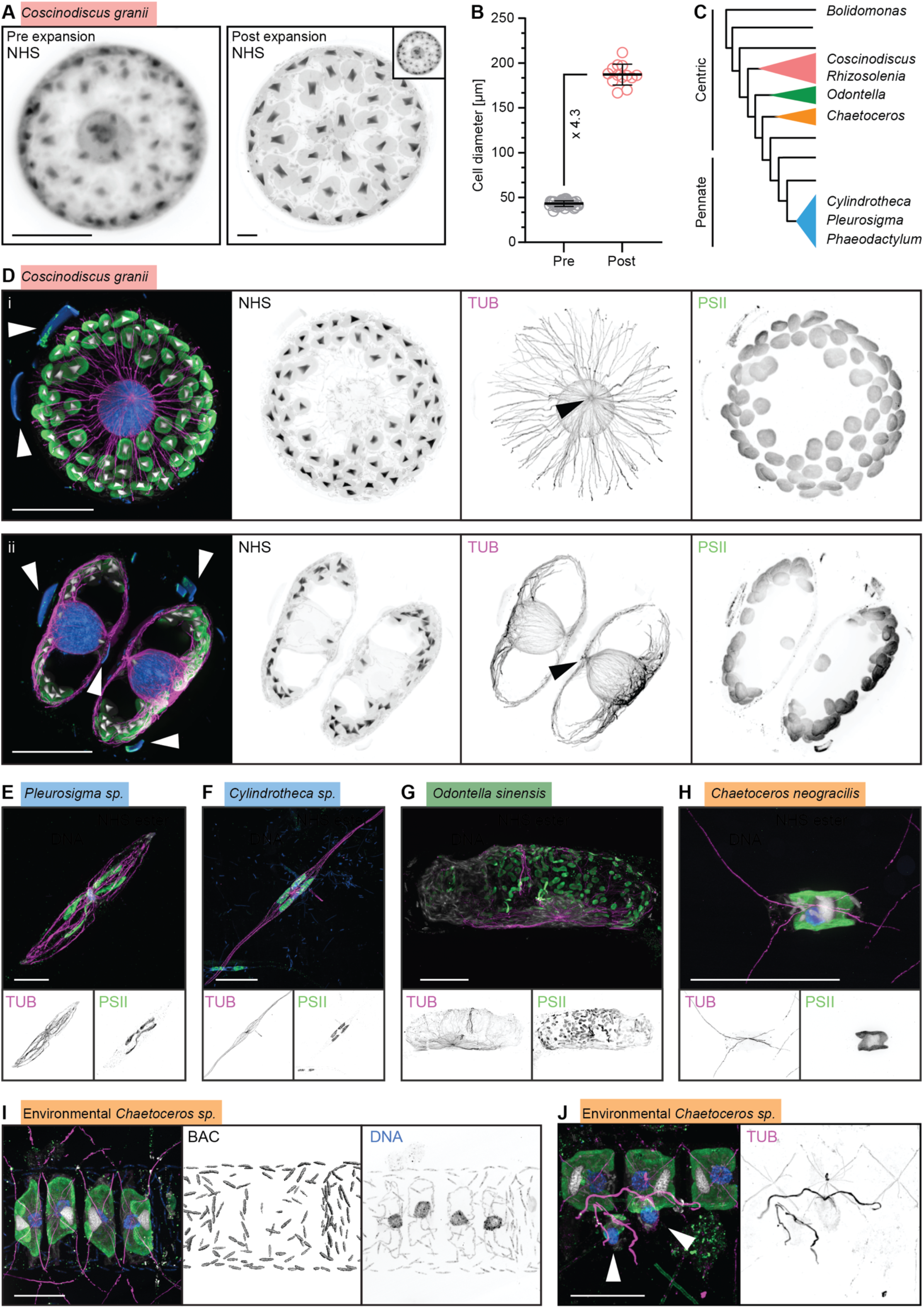
cryo-ExM enables high-resolution, three-dimensional imaging of cellular structures across several diatom species. (A) Maximum intensity projection showing NHS ester staining of a *Coscinodiscus granii* cell before and after expansion imaged using the same objective and microscope. The inset indicates the unadjusted pre-expansion size of the cell. Scale bars represent 20 µm (4.65 µm biological scale). (B) Measurements of the diameter of *C. granii* cells before and after expansion (n=37 and 13, respectively). The mean expansion factor was determined to be 4.3-fold. (C) Phylogenetic relation of diatom species used here indicated on the phylogram, adapted from (Medlin & and Kaczmarska, 2004). Clade colours are kept throughout the image. (D) Immunostaining for microtubules (magenta) and PSII combined with NHS ester pan-labelling (grey) and DAPI (blue) labelling the DNA and fragmented frustule (white arrow) of *C. granii*. Maximum intensity projections are shown spanning half a cell (i) and the central section (ii) allowing for a clear distinction of the MTOC (black arrow) and a microtubule cage spanning the nucleus in addition to the cortical bundles. NHS ester staining preferentially labels the triangular pyrenoid sitting roughly central in the manyfold chloroplasts which are found along the periphery and, in doublets, as single plastids in close proximity to the MTOC. Scale bars 100 µm (23.26 µm biological scale). (E - H) Exemplary maximum intensity projections of two pennate diatom cultures, *Pleurosigma sp.* (E) and *Cylindrotheca sp.* (F) as well as two more centric diatom cultures, *Odontella sinensis* (G) and *Chaetoceros neogracillis* (H). NHS ester (grey), microtubules (magenta and greyscale below), PSII (green and greyscale below), and DNA (blue) were stained for. Scale bars 50 µm (E, F, H), 100 µm (G) (Biological scale 13 µm in E, 18.5 µm in F, 28 µm in G, and 12 µm in H). (I-J) Environmental *Chaetoceros sp.* cells collected and fixed in Tallinn, Estonia, were observed in close association with (I) bacteria coating the outside of the frustule. Bacteria were stained with DAPI and readily segmented (BAC). (J) Another *Chaetoceros* cell was fixed in close association with two biflagellated cells (white arrows) containing distinct plastids. Scale bars 50 µm (12.5 µm biological scale).

The NHS ester labelling in *C. granii,* first allowed for an overall understanding of protein distribution in the cell (**Fig. 1D, NHS**). In particular, dense triangular shapes were identified as pyrenoids, specialized organelles that enhance carbon dioxide (CO_2_) fixation by concentrating RuBisCO, the protein that catalyses the reaction through which inorganic carbon enters the biosphere during photosynthesis (Ellis, 1979). The identity of the pyrenoid was further confirmed by direct staining of RuBisCO using a specific antibody (Rao et al., 2025) (**Supp. Fig. 2A**). While DAPI primarily stains for nucleic acids, unspecific staining was also detected on the fractured silica cell wall (**Fig. 1D, white arrows, Supp. Movie 1**). Vitrification likely creates microfractures in the frustule, facilitating expansion, additionally improving the preservation of many cellular features and epitopes leading to superior staining (Hinterndorfer et al., 2022). By using antibodies targeting tubulin, we aimed at visualising the microtubule network which lends structural support to cells and facilitates intracellular transport, frustule formation, and organelle positioning throughout the cell cycle. Within the same cell, we observed microtubule filaments and bundles along the periphery of the cell (**Fig. 1D, TUB, Supp. Movie 1**) as well as a cage-like structure of microtubules along the nucleus which form the cortical network, opposite of the polar microtubule organising centre (MTOC) proximal to the nucleus. Notably, diatom MTOCs appear to be acentriolar in line with previous literature (De Martino et al., 2009) featuring a prominent absence of microtubules-based cylindrical structure within their centre (**Fig. 1D, TUB, Supp. Movie 1**).

We then stained thylakoids, which are membranes within chloroplasts, by targeting PsbA or D1 protein, the core subunit of the PSII complex which provides electrons for all oxygenic photosynthesis to happen (Giordano et al., 2005), and thereafter named PSII. PSII fully surrounds the pyrenoids (**Fig. 1D, PSII**), and while most chloroplasts were localised regularly spaced along the periphery of the cell, they were excluded from the central area of both the epitheca and hypotheca (the upper and lower halves of the diatom silica cell wall), close to the nuclear interface (**Fig. 1D, PSII**). In some, but not all doublets, an exception was noted with individual chloroplast found at the MTOC of postmitotic cell doublets, close to the hypotheca, while the majority localises at the cell periphery of the epitheca (**Supp. Fig. 3**). This may represent a stage in the cell cycle ensuring even distribution of organelles during division. Differential intensity in PSII staining could be due to one side of the cell being closer to the coverslip, generating brighter signal due to light refraction with increasing depth (**Fig. 1D, PSII**).

To generalize the applicability of cryo-ExM to diatoms, we further expanded diverse cultures as well as environmental samples. Cultures of the pennate diatoms *Pleurosigma sp., Cylindrotheca sp*., the centric diatoms *Odontella sinensis* and *Chaetoceros neogracilis* and the three morphotypes of the model diatom *Phaeodactulym tricornutum* were successfully labelled (**Fig. 1C,E-H, Supp. Fig. 2,4**). These species diverged millions of years ago, with *Chaetoceros* having a more ancient origin (130 million years ago) compared to the relatively recent *Odontella* (35 million years ago) (Nakov et al., 2018) (**Fig. 1C**). Gel expansion was consistent across species of pennate and centric diatoms imaged here, however population level comparisons of cells pre- and post-expansion show slight variations in the level of expansion (**Supp. Table 1**) (Büttner et al., 2021). We then tested the applicability of cryo-ExM on diatoms taken from their natural environment. We took advantage of the pan-European TREC expedition and the Advanced Mobile Laboratory, a fully equipped molecular lab outfitted with cutting-edge instrumentation for in-depth biological analysis, in particular a plunge freezer for on-site vitrification. Planktonic communities collected and fixed in Tallinn (Estonia) and Plentzia (Spain) were expanded, stained, and imaged after having been stored in liquid nitrogen for several months. Remarkably, environmental samples expanded well (**Fig. 1I,J, Supp Fig. 1C-F**) showing that this state-of-the-art imaging approach can be relatively easily applied to complex environmental samples. On top, cryo-ExM can also successfully preserve cell-cell interactions that occur in environmental samples: bacteria were observed all along a *Chaetoceros* chain populating the extracellular surface (**Fig. 1I, Supp. Movie 2**), though the exact nature of the symbiosis - either mutualistic or parasitic - cannot be deciphered here. Surprisingly, bacteria seem to be organized along both longitudinal and lateral axes of the diatom chain, but not populating the setae, (**Supp. Fig. 5**), suggesting some form of specific colonization pattern, perhaps driven by fucose glycoconjugates as shown in other diatom species (Tran et al., 2023). Furthermore, small flagellated cells containing two chloroplasts and pyrenoids each were detected in close proximity to *Chaetoceros* chains from another species (**Fig. 1J, Supp. Fig. 1F**), recapitulating an interaction previously observed (Colin et al., 2017). Together, this demonstrates that cryo-ExM successfully overcomes diatom frustule barriers to achieve roughly 4-fold expansion across evolutionarily diverse species from both laboratory cultures and environmental samples, enabling detailed visualization of cellular structures while preserving ecological interactions.

### Systematic investigation of microtubule organisation across freely cycling diatom species

Diatoms are one of the earliest models for studying cell division due to their ease of culturing and their clear frustules (Cande & McDonald, 1985; Lauterborn, 1896; J. Pickett-Heaps, 1991; J. D. Pickett-Heaps & Tippit, 1978). Given the crucial role of microtubules in cell division and mitosis, both the organisation of microtubules and their roles in these functions have been explored across many diatom species. Previous descriptions of microtubule organisation in diatoms describe the localization of microtubules in interphase cells as extending from the MTOC near the nucleus towards the frustules (Tesson & Hildebrand, 2010; Tippit & Pickett-Heaps, 1977). It has been reported that as diatom cells enter mitosis, a polar complex (PC) appears outside the nucleus in pre-mitotic cells, a structure independent of the MTOC. The MTOC then disappears, and the PC later becomes the poles of the mitotic spindle (De Martino et al., 2009). After mitosis, the microtubules in a few diatom species have been shown to reorganize and expand, aiding in the positioning of silica deposition vesicles (SDVs) as a new valve forms (Tesson & Hildebrand, 2010). Furthermore, cryo-ET studies of *C. tenuissimus* (Mayzel et al., 2021) suggest that the microtubules can even extend into setae. Although actin was previously thought to be the cytoskeletal filament responsible for setae tip growth (J. D. Pickett-Heaps, 1998b), the presence of microtubules in the setae support more recent work suggesting that microtubules may also play a role in tip growth through positioning of silica deposition vesicles or may help transport other materials along the setae (Mayzel et al., 2021; J. D. Pickett-Heaps, 1998a; Tesson & Hildebrand, 2010). Despite the presence of these results, it is still difficult to make comparisons between species, and even between different cell cycle stages in the same species due to the variability in techniques and nomenclature. Additionally, there is a heavy bias in the literature towards pennate diatoms as one of the model diatom species most amenable currently to genetic manipulations is *P. tricornutum,* a pennate diatom. Here we use cryo-ExM to move past this barrier, and to highlight its potential as a tool for making qualitative and quantitative comparisons of the cytoskeleton throughout diatoms, even across pennate and centric species.

We compared interphase MTOC organisation across species spanning both centric (*C. neogracilis and C. granii*) and pennate (*Cylindrotheca sp. and E. clementina*) diatoms, and observed that the microtubules mirror the cell’s symmetry (**Fig. 2A, Supp. Fig. 6**). This is particularly striking in the centric *C. granii* and pennate *Cylindrotheca sp.* which display radial and bilateral symmetry respectively in both cell and microtubule organisation. We also observed a higher density of microtubules around the nucleus, as well as the localization of the MTOC on one side of the nucleus with microtubules extending from the MTOC towards the edges of the cell in all species we imaged (**Fig. 2A**). This suggests that diatoms share an interphase MTOC organisation despite their morphological diversity, and confirms previous results locating the MTOC as near the nucleus with microtubules extending towards the frustules. Furthermore, in the cultured *C. decipiens* and *C. neogracilis*, the microtubules extend to the corners and into the setae (**Fig. 2A,B**) as suggested in cryo-ET studies of *C. tenuissimus* (Mayzel et al., 2021), supporting the hypothesis that microtubules play a role in setae tip growth.

**Figure 2:**
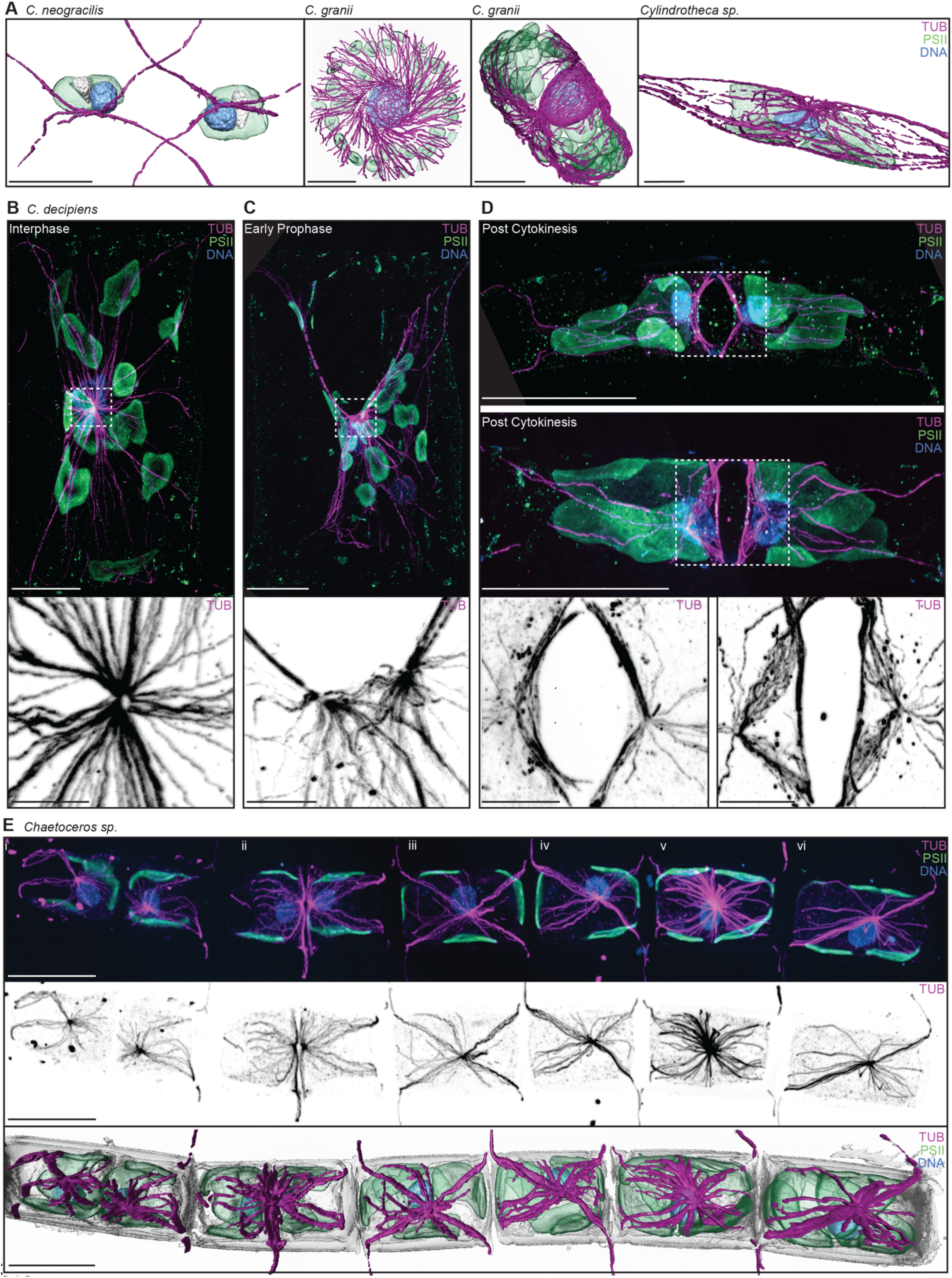
Characterization and identification of various cell cycle stages in both environmental and cultured samples. (A) 3D reconstructions of *C. neogracilis, C. granii* and *Cylindrotheca sp*. All reconstructions show segmented microtubules (magenta), chloroplasts (green) and nuclei (blue). The microtubules in each are localized to the side of the nucleus and extend to the edges of the cell. From left to right: expanded scale bars are 20 µm, 50 µm, 50 µm, 10 µm, and biological scale bars are 4.91 µm, 11.52 µm, 11.52 µm, 3.72 µm. (B-D) Characterization of different stages of the cell cycle in *C. decipiens.* (B) A cell in interphase presents the microtubules (magenta) generated from the MTOC towards the corner of the cell. The MTOC is located to the side of the nucleus (blue), and the chloroplasts (green) are distributed throughout the cell. Expanded scale bar 50 µm, biological scale bar 9.45 µm. The inset below of the microtubules in greyscale shows the ring-like structure of the MTOC. Expanded scale bar of the inset is 10 µm and the biological scale bar of the inset is 1.89 µm. (C) The next image is of a cell in early prophase. The microtubules now form two MTOCs, which the greyscale inset of microtubules is showing. The nucleus (blue) is located in the centre of the cell, and the chloroplasts (green) are still distributed throughout the cell. Expanded scale bar 50 µm, biological scale bar 9.45 µm for the top, composite image, and expanded scale bar 10 µm, biological scale bar 1.89 µm for the bottom greyscale image. (D) The top images are two cells which have just undergone cytokinesis. The MTOC is located near the plane of division in the top cell and left greyscale inset. In the bottom image and right inset, the MTOCs (purple and grey, respectively) are located further from the plane of division and closer to the centre of each of the daughter cells. The nucleus in both post-cytokinesis cells are still located near the plane of division and have not yet moved towards the centre of the cell. Expanded scale bar 50 µm, biological scale bar 13.05 µm for the composite panel and expanded scale bar 10 µm, biological scale bar 2.62 µm for tubulin-only greyscale panel. (E) The top panel is an environmentally sampled chain from Plentzia, Spain. The nucleus (blue), chloroplasts (green) and microtubules (magenta) are all useful for identifying which stages of the cell cycle each cell in the chain is in. The microtubules are shown in the middle panel in grey scale to highlight the MTOCs and reach of the microtubules in the different cells. The bottom panel shows a 3D reconstruction of the same chain. The two left cells in the chain (i-ii) have recently divided, with the second cell (ii) more recently having undergone cytokinesis. The cells on the right (iii-vi) are all in interphase. Expanded scale bar 50 µm, biological scale bar 12.63 µm.

Cryo-ExM also allows us to qualitatively and quantitatively make comparisons across different stages of the cell cycle. We focus on one species, *C. decipiens,* a centric diatom whose mitotic spindle had been previously observed (J. D. Pickett-Heaps, 1998a), but whose microtubule organisation throughout the other stages of the cell cycle has not previously been characterized (Darley & Volcani, 1971). In the interphase of *C. decipiens* cells, microtubules originate from a ring-shaped microtubule organising center (MTOC) near the nucleus (**Fig. 2B**). This is similar both to the literature and to our observations of the MTOC in interphase of *C. granii* and *Cylindrotheca sp.* cells. As cells progress from interphase to early prophase, two MTOCs become visible in *C. decipiens* (**Fig. 2C**). Just after cytokinesis, the two MTOCs remain at the plane of division, and later are observed closer to the centre of the cell (**Fig. 2D**). The localization of the MTOCs post-cytokinesis to the plane of division, where the new frustule will have to form, further implicates microtubules in the valve formation process (Tesson & Hildebrand, 2010). Throughout all these stages, the MTOCs are consistently localized to one side of the nucleus (**Fig. 2B-D**). These results vary from previous literature, which has shown that the MTOC disappears and reforms after mitosis, and has not previously suggested that the MTOC duplicates prior to mitosis (De Martino et al., 2009). Although we cannot discount that the MTOC may still disappear and reform after mitosis, the presence of two MTOCs prior to mitosis and near the plane of division post cytokinesis suggests that this may not be the case, however further studies of the other stages of mitosis both in *C. decipiens* and in other diatom species would be needed to further answer this question.

Characterizing the stages of the cell cycle through microtubule organisation in *C. decipiens* allows us to understand how cell division varies not only across species or across cell cycle stages, but across cells in a community. Diatoms in culture are not naturally synchronised, although they can be synchronised through silicon starvation or light starvation (Darley & Volcani, 1971). However, diatoms in chains have been shown to have synchronized divisions(J. D. Pickett-Heaps, 1998a). These contradicting results can be further explored, even in cells sampled from natural populations, using the higher throughput and more systematic approach of cryo-ExM. We established this by expanding an environmentally sampled *Chaetoceros* chain from Plentzia, Spain (**Fig. 2E, Supp. Movie 3,4**), which contains, from right to left, four cells in interphase (iii-vi), a cell which has recently undergone mitosis and has just undergone cytokinesis (ii), and a cell which has recently divided and physically separated the MTOCs of the daughter cells towards the centre of each cell (i). The microtubules of each of these cells extend to the corner of the cell and into the setae when the setae are present, as observed in cultures. The presence of cells which have recently divided next to each other suggests that division in chains, especially those from natural samples as opposed to cultures, may not be as synchronised as was previously reported. However, this does not rule out the effect of abiotic factors on the level of synchrony within a chain or within a population, as was suggested by the synchronised culture studies. A larger scale study of microtubule organisation on natural populations of cells in an environment with changing abiotic conditions could answer this question, and is now possible given cryo-ExM. The ability to now target specific structures in order to characterize and identify different stages of the cell cycle is extremely valuable, especially in natural samples. Identifying stages in the cell cycle in environmental cells provides a method to understand the molecular mechanisms behind synchronization, timescales of division, and diversity of cellular division apparatuses across species in their natural habitats.

### Plasticity of chloroplast and pyrenoid architectures within and across diatom species

The chloroplast is the organelle responsible for fine tuning key essential processes in diatoms such as photosynthesis and photoprotection, which occur in the thylakoid membranes, as well as carbon fixation and CO_2_ Concentrating Mechanisms (CCMs), which takes place in the pyrenoid. While chloroplasts and pyrenoids ultrastructure may reflect certain ecological niches and vary among species (Barrett et al., 2021; Bedoshvili et al., 2009), they may also provide clues on the physiological state of the cells and how the surrounding environment affects them through reduced or enhanced CO₂ availability or luminous stress (Broderson et al., 2024; Polukhina et al., 2016; Wood et al., 2019). Studying functional plasticity in fluctuating environments can be extremely challenging especially in non-model organisms. So far, structural visualization of both photosynthetic membranes and pyrenoids was possible through high-resolution albeit low-throughput electron microscopy (EM), Focus-Ions Beam Scanning Electron Microscopy (FIB-SEM) and cryo-electron tomography (cryo-ET) and a direct morphological investigation, using light microscopy, had not been attempted in diatoms. Taking advantage of cryo-ExM’s improved spatial resolution, we acquired z-stack from PSII and NHS, segmenting chloroplasts and pyrenoids volumes in both cultured and environmental species, centric and pennate diatoms to better capture phenotypic plasticity of their photosynthetic machinery.

Our comparative analysis of chloroplasts and pyrenoids morphologies across species shows that characteristic shapes are unique to each diatom. In the large centric diatom *C. granii*, multiple oval-shaped chloroplasts surround the entire cell and the pyrenoids possess a pyramidal shape (**Fig. 3A, Supp. Movie 1**). The environmental *Chaetoceros sp*., displayed four complex lobe-shaped chloroplasts per cell and an equal number of flattened, rectangular pyrenoids (**Fig. 3B, Supp. Movie 4**). In the cultured centric diatom *C. neogracilis*, we observed one round chloroplast per cell and a single rounded pyrenoid with often a pointed edge (**Fig. 3C, Supp. Movie 5**). Finally, the raphid pennate diatom *Cylindrotheca sp*. displayed two large elongated chloroplasts and two pyrenoids with a lenticular shape (**Fig. 3D**). We further quantified surface/volume ratios in chloroplasts and pyrenoids to investigate individual shape distribution and variability among different species (**Fig. 3E**). Despite their differences in shape, the pyrenoid surface/volumes ratios were surprisingly similar in *C. granii* cultures and the *Chaetoceros sp*. cryo-fixed in Plentzia, Spain (1.22 ± 0.1; 0.93 ± 0.14, respectively). In the other two cultures, *C. neogracilis* and *Cylindrotheca sp*., the average ratio was higher (1.47± 0.12; 1.96 ± 0.06, respectively). Likewise, the pyrenoids/chloroplasts ratio (%) was preserved in all diatoms investigated (**Fig. 3F**). In two different *C. granii* cells, the pyrenoid/chloroplast ratio was 5.9% (n=27) and 6.5% (n=17); in two two-celled chains of *C. neogracilis* the ratio was 11.2% (n=2) and 11.7% (n=2), in two different *Cylindrotheca sp*. cells the ratio was 11.3% (n=2) and 7.4% (n=2). In the environmental *Chaetoceros sp.* chain composed of six cells, the pyrenoid/chloroplast ratio was 9.1% (n=24). In the case of *C. granii*, lower values are possibly compensated by the larger number of those organelles in the cell.

**Figure 3:**
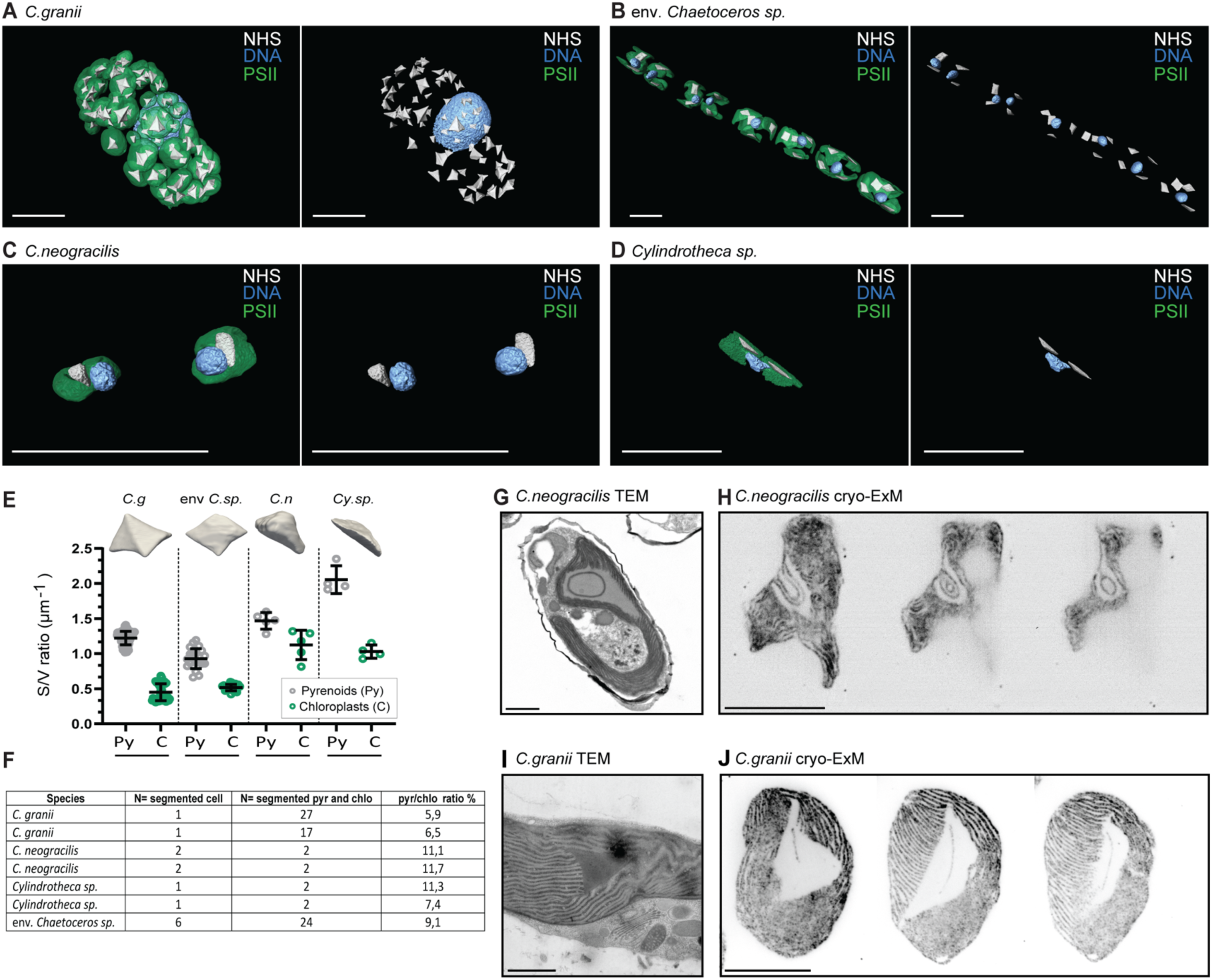
Phenotypic plasticity of the photosynthetic machinery (chloroplast, pyrenoid and Pyrenoid Penetrating Thylakoids). (A-D) 3D segmentation of pyrenoids, chloroplasts and nuclei in (A) the large centric diatom *C.granii*, expanded scale bar 50 µm, biological scale bar 11,52 µm; (B) the env. *Chaetoceros sp.* isolated in Plentzia, expanded scale bar 50 µm, biological scale bar 12,5 µm; (C) the *C. neogracilis* culture, expanded *s*cale bar 50 µm, biological scale bar 12,29 µm; (D) the raphid pennate culture *Cylindrotheca* sp., expanded scale bar 50 µm, biological scale bar 18,59 µm. (E) Surface/volume ratio quantification in chloroplasts and pyrenoids. On the upper part an illustration of the pyrenoid shape is added. Grey and green dots represent pyrenoid and chloroplast surface/volume ratio respectively for *C. granii* (1.22 ± 0.1; n=89, in 2 segmented cells), the env. *Chaetoceros sp.* (0.93 ± 0.14, n=22, in 6 segmented cells forming one chain)*, C.neogracilis* (1.47± 0.12; n=5, in 5 segmented cells) and *Cylindrotheca sp* (1.96 ± 0.06, n=2, in 2 segmented cells). (F) Table highlighting the segmented species, where each row is a separate acquisition; number of segmented cells in the case of chain forming species; number of segmented pyrenoids and chloroplasts in each cell and finally pyrenoid/chloroplast volume ratio. (G) *C. neogracilis* TEM section showing the pyrenoid with a round PPTs ring inside. Scale 1 µm (H) Montage of 3 slides with increasing z-depth showing *C. neogracilis* thylakoids and PPTs in cryo-ExM. Expanded scale bar 10 µm, biological scale bar 2,5 µm; (I) *C. granii* TEM section showing a triangular side pyrenoid. Scale 1 µm. (J) Montage of 3 slides with increasing z-depth showning *C. granii* thylakoids and PPTs in cryo-ExM. Expanded scale bar 10 µm, biological scale bar 2,5 µm.

Diatom pyrenoids conserve some robust characteristics among different diatom species (Bedoshvili et al., 2009), one fascinating example being the Pyrenoid Penetrating Thylakoids (PPTs). They enable tight spatial and temporal control of CO_2_ delivery to the RuBisCO, via Carbonic anhydrases (CAHs) thus facilitating high-efficiency CO_2_ concentrating mechanisms (Matsuda et al., 2011). However, their direct role and dynamics in CO_2_ Concentrating Mechanisms (CCMs) have remained largely understudied and have, to date, remained confined to laboratory model systems. We therefore compared PPTs in *C. neogracilis* and *C. granii* using both electron and expansion microscopy. In *C. neogracilis*, the PPTs follow the round shape of the pyrenoid in both imaging modalities (**Fig. 3G,H**). In *C. granii*, the electron microscopy section showed a triangular side pyrenoid confirming the pyramidal shapes seen in expansion (**Fig. 3I**), with PPTs perpendicular to the pyrenoid (**Fig. 3J**). The concordance between these two complementary approaches enabled us to better understand PPTs in natural samples from Tallinn, where *Chaetoceros* PPTs show a triangular V-shape (**Supp. Fig. 7A**).

Overall, these results provide valuable qualitative and quantitative insights on photosynthetic diversity, indicating that cryo-ExM could now be used to investigate light acclimation and CCMs strategies in diatoms. We can indeed speculate that in CO_2_-rich environments some species modulate diffusion barrier structures (i.e., by reducing they surface/volume ratio; pyrenoids in rounded shape), while species adapted to unpredictable environments may opt for more flattened structures for maximized CCMs thus reducing CO_2_ diffusion.

## Perspective

Here, we provide in depth validation of cryo-ExM in diatoms, showcasing its applicability across several phylogenetically distant species as well as canonical model systems, in the lab and in the field. We illustrate its potential for comparative cell biology in diatoms by targeting two key proteins, tubulin and PsbA as well as pyrenoids using NHS and show how both microtubule and chloroplast architectures can be rapidly profiled across cultured and environmental cells, including in symbioses. We characterize microtubule organisation across the cell cycle of individual species, and across cells within a population, and reveal similarities in interphase microtubule organisation across many species of diatoms which includes both pennate and centric morphologies. Furthermore, the comparative analysis of chloroplasts and pyrenoids morphologies across species provides qualitative and quantitative insights on photosynthetic diversity, suggesting that photosynthetic architectures are peculiar to each diatom.

Based on this study, many exciting perspectives can be outlined to advance cell biology in diatoms. Recently, two independent groups identified the PyShell (Nam et al., 2024; Shimakawa et al., 2024), a protein shell that encases the diatom pyrenoid and plays a critical role in defining the pyrenoid’s shape, and used genetically fused candidate proteins to fluorescent reporters in *T. pseudonana*. This work presents promising results, but is constrained by the necessity of genetic transformation which is limited in diatoms. Staining for proposed PyShell proteins with generated antibodies may now be possible across a range of diatoms and immediately provide higher resolution data to better understand pyrenoid shape-shifting in different diatom species and under different environmental conditions. Localizing proteins involved in silica cell wall formation such as silicic acid transporters (SITs), localized to the plasma membrane and potentially other intracellular membranes, could improve our understanding of the uptake of silicic acid necessary for cell wall synthesis. Additionally, silaffins are central to silica polymerization and are embedded in specific regions of the biosilica, such as the valve region. Precise localization of silaffins like Silaffin-3 (tpSil3) in *T. pseudonana* revealed distinct accumulation areas within the biosilica, crucial for understanding frustule morphogenesis, and could here be visualized across different species (Gröger et al., 2016). Beyond protein labelling, coupling of cryo-ExM with mRNA and rRNA *in situ* hybridization could open opportunities for further genotype to phenotype mapping.

Beyond its potential in unravelling key aspects of diatom cell biology, cryo-ExM could contribute to shedding more light on diatom ecology and evolution. For instance, the localization of proteins involved in growth limiting iron uptake such as phyto transferrin (ISIP2A) could be investigated in cells with specific morphological features (such as *Chaetoceros* setae) or diatom biofilms to see if some regions specialise in nutrient uptake more than others. Triose phosphate translocators (TPTs) are localized in specific membranes of the complex plastid envelope, mediating carbohydrate exchange between the endosymbiont (plastid) and the host cytosol. Their correct localization is fundamental for establishing and maintaining the metabolic connection between host and endosymbiont, a key evolutionary innovation in diatoms that has so far been investigated in a handful of species (Moog et al., 2015). Antioxidant proteins such as superoxide dismutase (SOD) and glutathione S-transferase (GST) need to be localized near sites of reactive oxygen species (ROS) production, such as chloroplasts or mitochondria, to effectively mitigate oxidative stress induced by abiotic changes such as salinity and could be studied within euryhaline diatoms such as *Cyclotella*, that can tolerate and grow across a wide range of habitats, from nearly freshwater to hypersaline conditions (Downey et al., 2023). Furthermore, cryo-ExM may be particularly relevant for the study of overlooked nano diatoms (Leblanc et al., 2018), that sometimes reach less than 2µm in diameter but play an important role in marine ecosystems.

With this study, we hope to generate new opportunities in diatom cell biology, by enabling detailed spatial resolution of cellular structures across diverse diatom species in both laboratory and natural environments. As our climate changes and oceans transform, understanding the cellular foundations of diatom adaptability becomes increasingly urgent. We invite researchers to rediscover diatoms as powerful model systems at the intersection of cell biology and ecology, unlocking secrets that may guide our understanding of resilience in a changing world.

## Data availability

All data is available in the following s3 storage bucket: https://console.s3.embl.de/browser/diatomultrastructuraldiversity and is available with the access key eCQoR9HTs7hvLYJ0qHm9 and the secret key aY3u2zcXuDEaI92eqS67xwbxjyNylqY7P8kQVkWq. The data can be accessed using a miniIO client or any other s3 compatible client.

## Acknowledgements

We thank Chris Bowler, Paul Guichard, Virginie Hamel, Ben Engel and members of the Vincent and Dey Groups for their feedback on the manuscript. Special thanks to the TREC Consortium members, the TREC Expedition team, the TREC core partners EMBL, the Tara EUROPA team, the Tara Ocean Foundation and EMBRC and in particular our local partners Sirje Sildever from TalTec, Estonia, Ibon Cancio from UvP, Spain and the captains for hosting and guidance on sampling sites. We thank Sophie Baars, Chandni Bhickta and Clemence Anne Saint-Donat for sampling plankton during the TREC Expedition. We thank Alessandra Mussi for guidance on diatom phylogeny. We acknowledge services provided by the Mobile Laboratories Facility of the European Molecular Biology Laboratory.

## Author contributions

S.F, F.M, E.F, K.M, S.G, I.R-B and V.O developed methodology, conducted formal analysis, curated and visualised data. F.V, G.D, O.D, T.B, S.Z, M.O, M.L, M.S-O provided resources. F.V, G.D, Y.S, O.D, E.Y acquired funding. F.V and G.D supervised the study. S.F, F.M, E.F, G.D and F.V conceptualized the study and wrote the original draft. All authors reviewed and edited the manuscript.

## Material and Methods

### Algal cultivation

All algal monoclonal cultures were grown in 25 mL single-use flasks at low light intensity of 40 µmol photon m^-2^. S^-1^ and a 12/12-hour light/dark photoperiod at 15°C (incubators) and refreshed weekly using F/2 medium enriched with silica. The laboratory strains used in this study are *Coscinodiscus granii* RCC7046, *Odontella sinensis* RCC7047, *Phaeodatylum tricornutum* CCMP2561, *Chaetoceros neogracilis* TREC16 (Tallin, Estonia), *Pleurosigma sp* TREC112 (Barcelona, Spain), *Cylindrotheca sp* TREC24 (Kristineberg, Sweden), *Rhizosolenia setigara* TREC2 (Roscoff, France).

The TREC strains were isolated during the TREC expedition thanks to the Advanced Mobile Laboratory. Briefly, plankton samples were taken daily and single planktonic cells were isolated using a COPAS Vision 500 (Union Biometrica) in 96 well flat bottom culture plates filled with 200 µL of F/50 media. After shipping to EMBL, cells were pre-screened using the microscope Olympus/Evident (CKX53). Interesting strains were picked and transferred with a 200µl pipette into a 24-well plate well containing 500µl F/2 (Guillard′s (F/2) Marine Water Enrichment Solution REF:G0154, Merck) supplemented with silica (F/2++). After 5-10 days cells were checked again and 500µl of good candidates were transferred to a 6-well plate containing 2 ml of F/2 ++ media. Eventually the strains were transferred again to a 25 ml culture flask with a filter cap (Nunc™ EasYFlask™ Cell Culture Thermo fisher 156367) containing 20 ml of F/2++. In those flasks, cells were checked regularly and refreshed roughly every 20 days. All cells were placed into an incubator (Percival SE-41P4) with adjusted light intensity (100 µEinstein – 150 µEinstein) and temperature (15°C – 20°C) on a 12/12 day and night cycle. Salinity of medium and incubator settings were adapted according to the conditions at the TREC stop where cells were isolated.

### Sample fixation of lab and field samples

#### Sampling

Lab cultures were harvested at exponential phase after the onset of the light period. Environmental samples were collected during the TREC expedition on 04.10.2023 at Billano Ira, Plentzia, Spain (43.439251, -2.9364519) and on 22.06.2023 at Tallinn, Estonia (59.44944, 24.78777) using 10 µm nets with content being filtered to 100 µm and 40 µm respectively. Water was maintained at sea temperature until further processing.

#### Fixation

Cryo-fixation and cryo-substitution were performed following previously published cryo-ExM protocol (Laporte et al., 2022), with minor modifications. Laboratory cultures and field samples were concentrated using mild vacuum filtration onto 1.2 µm mixed cellulose ester when required and mounted onto 12 mm coverslips coated with Poly-L-Lysine and/or CellTak. After letting cells settle and attach, residual supernatant was carefully removed and blotted off using tissue paper before rapidly plunging the coverslip into liquid ethane or a mix of ethane-propane using a manual plunge freezing system (RHOST LLC). Coverslips were placed into home-made metal coverslips for cryo storage and kept at liquid nitrogen temperature until further processing. For freeze substituting samples, coverslips were placed into tubes containing frozen acetone containing 0.5% (v/v) formaldehyde and 0.025% (v/v) glutaraldehyde, fully submerged in dry ice and incubated overnight. The following day, dry ice was gradually removed over the course of ∼6 hours to slowly increase the temperature. In order to rehydrate the samples, coverslips were moved into ethanol once at room temperature and washed with increasing dilutions of ethanol (100%, 95%, 75%, 50%, 0% (v/v) ethanol in water) for 5 min each. Cells that detached during the process (most commonly in the acetone or 100% ethanol steps) were collected, processed in the same fashion and remounted onto fresh coverslips when required. All samples were stored at 4°C in PBS until further use.

### U-ExM workflow

U-ExM was performed following previously published protocol (Gambarotto et al., 2019). Fixed and rehydrated samples were incubated with 1% acrylamide and 0.7% formaldehyde in PBS overnight at 37°C. Subsequently, coverslips were inverted onto droplets of monomer solution [10% (v/v) acrylamide, 19% (v/v) sodium acrylate, 0.1% (v/v) N,N′- Methylenebisacrylamide, 0.5% (v/v) APS, and 0.5% (v/v) TEMED in PBS] on ice and incubate for 5 min on ice. Gels were polymerised fully by incubating them in humid chambers for 45 min – 1.5 h at 37°C. Samples were gently detached from coverslips and either moved into a preheated denaturation buffer or transferred into a 6-well plate filled with denaturation buffer in a rotating shaker with moderate speed for 15-20 min. Gels will separate from coverslips. Samples were denatured at 95°C for 1.5 h in denaturation buffer [200 mM SDS, 200 mM NaCl, 50 mM Tris-HCl adjusted to pH 9.0], washed once with PBS to remove residual buffer and fully expanded by incubating in ddH2O three times for 30 min each. Gel diameters were measured as an approximation of the expansion factor, cut into pieces when required and shrunken again by washing three times with PBS for 15 min each. Antibody stainings were performed in 4% BSA (w/v), 0.2% - 0.05 % (v/v) Tween-20 in PBS. The following primary antibodies were used: mouse monoclonal anti-tubulin 12G10 (AB_1157911, IgG, DHSB USA, 1:1000), rabbit polyclonal anti-PsbA-D1 protein of Photosystem II (PS2; IgG, Agrisera Sweden; AS05084, 1:1000) used as proxy for thylakoid membranes ultrastructure, and rabbit polyclonal anti-RbcL (Agrisera Sweden ref AS03037, 1:1000). Primary antibodies were used at indicated dilutions and incubated at 37°C overnight. Secondary antibodies were anti-mouse (IgG) and anti-rabbit (IgG) conjugated with Alexa Fluor 555 (Thermo Fisher Scientific, A32727 & A11011; 1:1000) or Alexa Fluor 488 (Thermo Fisher Scientific, A11034 & A11001; 1:1000). Secondary antibodies were incubated at 37°C for 4 h. Samples were washed three times with PBS 0.1% Tween-20 for 10 min each at room temperature. NHS-ester (at 1 µg/ml) coupled to AlexaFluor 647 (Invitrogen, A37573) and DAPI (at 5 µg/ml) in PBS were incubated overnight at 4°C. After staining, gels were re-expanded in ddH2O by washing them three times for 30 min each before acquisition.

### Image acquisition

Fully expanded gels were mounted onto poly-L-Lysine coated glass bottom dishes with the side of the gel containing cell facing the glass bottom. Gels were kept moist throughout the acquisition to prevent shrinkage. Microscopy was performed using a Nikon-CSU-W1 SORA spinning disk confocal microscope with water immersion objectives (Apo LWD 40x 751 WI Lambda-S/1.15, 0.60 and SR P-Apochromat IR AC 60x WI/1.27) and a CSUW1-TS 2D 752 50/SoRa disk. In some cases, an additional 4x SoRa disk was employed to further increase magnification. Overview acquisitions were taken using a P-Apo Lambda S 10X/ 0.45/ WD 4,0 /air objective.

### Image analysis

Images were processed with FIJI (ImageJ). Images were processed to adjust brightness linearly and analysed using FIJI. Stacks were flattened and are shown where indicated as average or maximum projections. The expansion factor was calculated as described in (Gambarotto et al., 2019). In addition, direct measurements of the cells pre-expanded and post-expanded in gel were performed. Two different diatom species were measured: *C. granii* cells n=13 using the diameter (expansion factor 4.3) and C. *neogracilis* cells n=30 using both length and width (expansion factor 3.4 length; width 4.1). All the 3D segmentation, visualisation and volumetric analysis were made in Amira (v.2024.2; ThermoFisher Scientific). Graphs were prepared in GraphPad Prism 10 (GraphPad Software, Boston, Massachusetts USA, www.graphpad.com). Animations were made with Imaris (Oxford Instruments).

### Confocal microscopy of cultures

Live cell imaging was performed using a Zeiss LSM900 confocal laser scanning microscope equipped with a 40x/1.4 NA Oil DIC M27 objective lens. Samples were mounted on poly-L-Lysine coated glass bottom dishes. A 488 nm laser was employed for chlorophyll fluorescence imaging and Z-stack images were acquired with optimal step size to ensure proper 3D reconstruction. Post-acquisition image processing, including brightness/contrast adjustments and 3D rendering, was performed using Zeiss ZEN Blue (v3.5) software.

### Transmission electron microscopy of C. neogracilis TREC16

#### Chemical fixation

In order to preserve the delicate frustule in *C. neogracilis* an optimized protocol was applied. Cells were fixed overnight at 4°C in 2% Paraformaldehyde (EMS 15710) with 0.2% Glutaraldehyde (Polysciences 00216) and 4% sucrose (Sigma 84097) in 0.1M PHEM buffer. After 4x rinsing in 0.1M PHEM buffer cells were postfixed in 0.1% OSO4 (EMS 19150) in 0.1M PHEM buffer for 2h at 4°C. Cells were rinsed in dH2O and dehydrated in an increasing concentration of ethanol followed by a stepwise infiltration in Spurr resin diluted in ethanol 100% (Sigma EM300-1KT, 33%-66%-4x100%). Resin embedded cells were then polymerized at 65°C for 48 h.

#### Ultramicrotomy and imaging

Ultrathin sections (80 nm) of the resin embedded *C. neogracilis* cells were obtained with an ultramicrotome (Leica EM UC7) and a 35° diamond knife (Diatome) and collected on 3 mm copper slot grids (Agar Scientific) coated with 1% formvar in chloroform. Sections on grids were post-stained in 1% Uranyl-acetate in dH2O for 20 min. followed by Reynolds Lead-citrate for 1 min. The sections were examined and imaged with a transmission electron microscope (TEM, JEOL1400 plus).

### Transmission electron microscopy of C. granii

#### High Pressure Freezing (HPF)

*C.granii* cells were transferred with a tip to an “A” gold-copper carrier (Leica Microsystems) soaked in 1-hexadecene (Merck). The excess of sea water/18exadecane was removed with absorbent paper points (Gapadent). Then an “B” aluminum carrier (Leica Microsystems) was placed on top with the flat side facing the cells. Subsequently, cells were cryofixed with a HPM010 machine (ABRA Fluid AG) and stored in LN2 until their further processing.

#### Freeze Substitution (FS)

Frozen *C. granii* were freeze substituted following the protocol of Stigloher and colleagues (Stigloher et al., 2011) with some modifications. Briefly, cells were incubated in 0.1% tannic acid and 0.5% glutaraldehyde in acetone for 92 h at 90°C in an automatic freeze substitution machine (Leica EM AFS). Cells were then washed five times with anhydrous acetone. Subsequently cells were incubated with a 2% OsO4 solution in acetone for 32 h at 90°C. The temperature was then raised over the course of 14 h to -20°C and kept for 16 h. Next, the temperature was raised over 4 h to 4°C and the OsO4 solution was removed by washing five times with anhydrous acetone. After warming up to room temperature (20°C), cells were infiltrated with 25%, 50% and 75% epoxy resin (EPON 812, Serva) in acetone (without the accelerator), 2 h each. After a 100% epoxy resin infiltration overnight, cells were embedded in a 100% fresh solution of epoxy resin for 68 h (both without the accelerator). Then cells were incubated three times (3 h/each) and once (overnight) with 100% epoxy resin (containing the accelerator, DMP-30). Resin embedded cells were then polymerized at 60°C for 48 h.

#### Ultramicrotomy and imaging

Ultrathin sections (70 nm) of the resin embedded *C. granii* were obtained with an ultramicrotome (Leica EM UC7) and a 35° diamond knife (Diatome) and collected on 3 mm copper slot grids (Agar Scientific) coated with 1% formvar in chloroform. The sections were examined and imaged with a transmission electron microscope (TEM, JEM 2100+, JEOL).

## Supplementary Figures

**Supplementary Figure 1:**
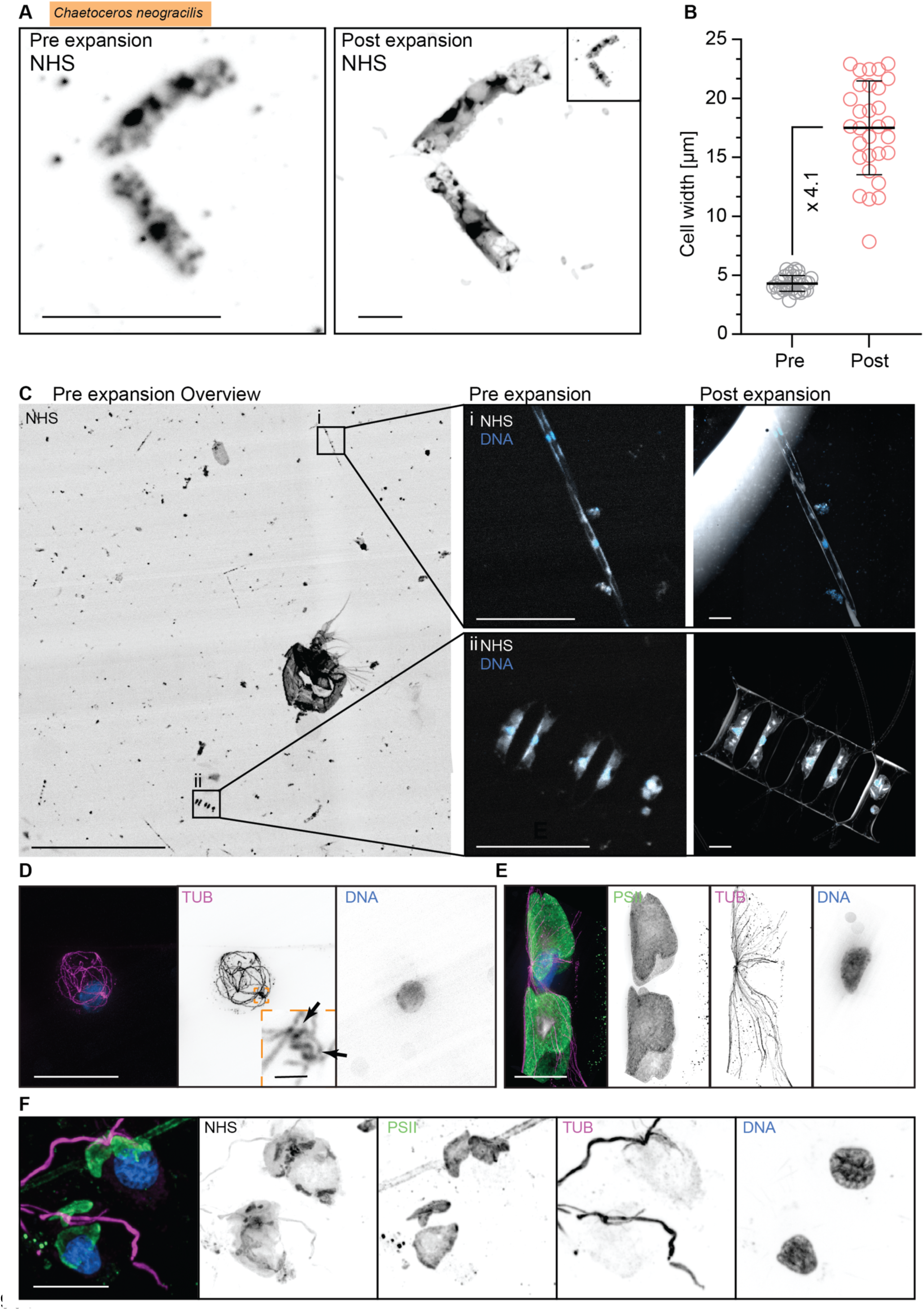
(A) Maximum intensity projection showing NHS ester staining of a *C. neogracilis* cell before and after expansion imaged using the same objective and microscope. The inset indicates the unadjusted pre-expansion size of the cell. Scale bars 20 µm (B) Measurements of the width of *C. neogracilis* cells before and after expansion (n= 30, respectively). The mean expansion factor was determined to be 4.1-fold. (C) Overview of a coverslip collected and fixed in Plentzia, Spain shortly after collection and stained with NHS ester. Several different species and cells including a fragmented copepod can be distinguished. The scale bar represents 500 µm and zoom in on two example cells from C shown with NHS ester and DAPI staining before and after expansion. Note that an air bubble introduced a large halo effect in NHS ester staining in the top right. Scale bars represent 50 µm. (D & E) Higher magnification acquisition of features from C showing two distinct areas within a chain of *Chaetoceros sp.*, with (D) showing a small spherical cell devoid of PSII staining but showing a elaborate microtubule (magenta) network and centriolar MTOC (inset and arrows, scale bar 2 µm). The neighbouring cell in the chain in (E) presents an expected morphology with a polar, nuclear (blue) associated centriole-less MTOC, lobes of plastids (green) with central pyrenoids (grey). The scale bars represent 20 µm. (F) Close up of microflagellates from Fig. 1J show distinct nuclei, two chloroplasts per cell, with discernible pyrenoids. Scale bar 20 µm.

**Supplementary Figure 2:**
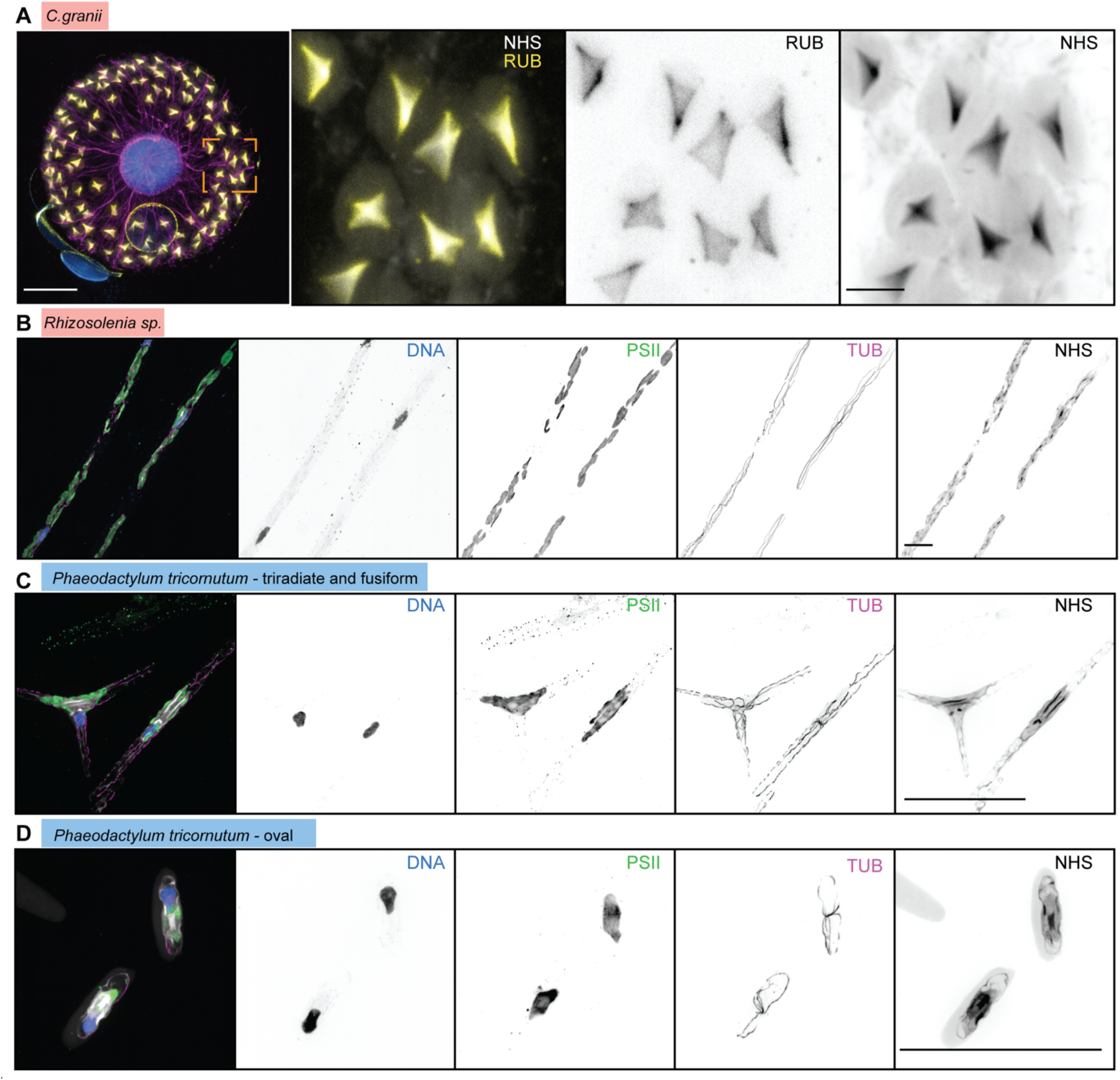
Rubisco staining and expanded pennates. (A) Left panel *Coscinodiscus granii* overview, expanded scale bar 50 µm, biological scale bar 11,52 µm. The three right panels zoom into the orange square with merged RuBisCo and NHS, RuBisCo, NHS ester, expanded scale bar 10 µm, biological scale bar 2,30 µm. (B) *Rhizosolenia sp.* Merge, DNA, PSII, tubulin, NHS ester, expanded scale bar 50 µm, biological scale bar 16,29 µm. (C) *P. tricornutum* triradiate and fusiform morphotype. Merge, DNA, PSII, tubulin, NHS ester, expanded scale bar 50 µm, biological scale bar 26,74 µm. (D) *P. tricornutum* oval morphotype. Merge, DNA, PSII, tubulin, NHS ester, expanded scale bar 50 µm, biological scale bar 26,74 µm.

**Supplementary Figure 3:**
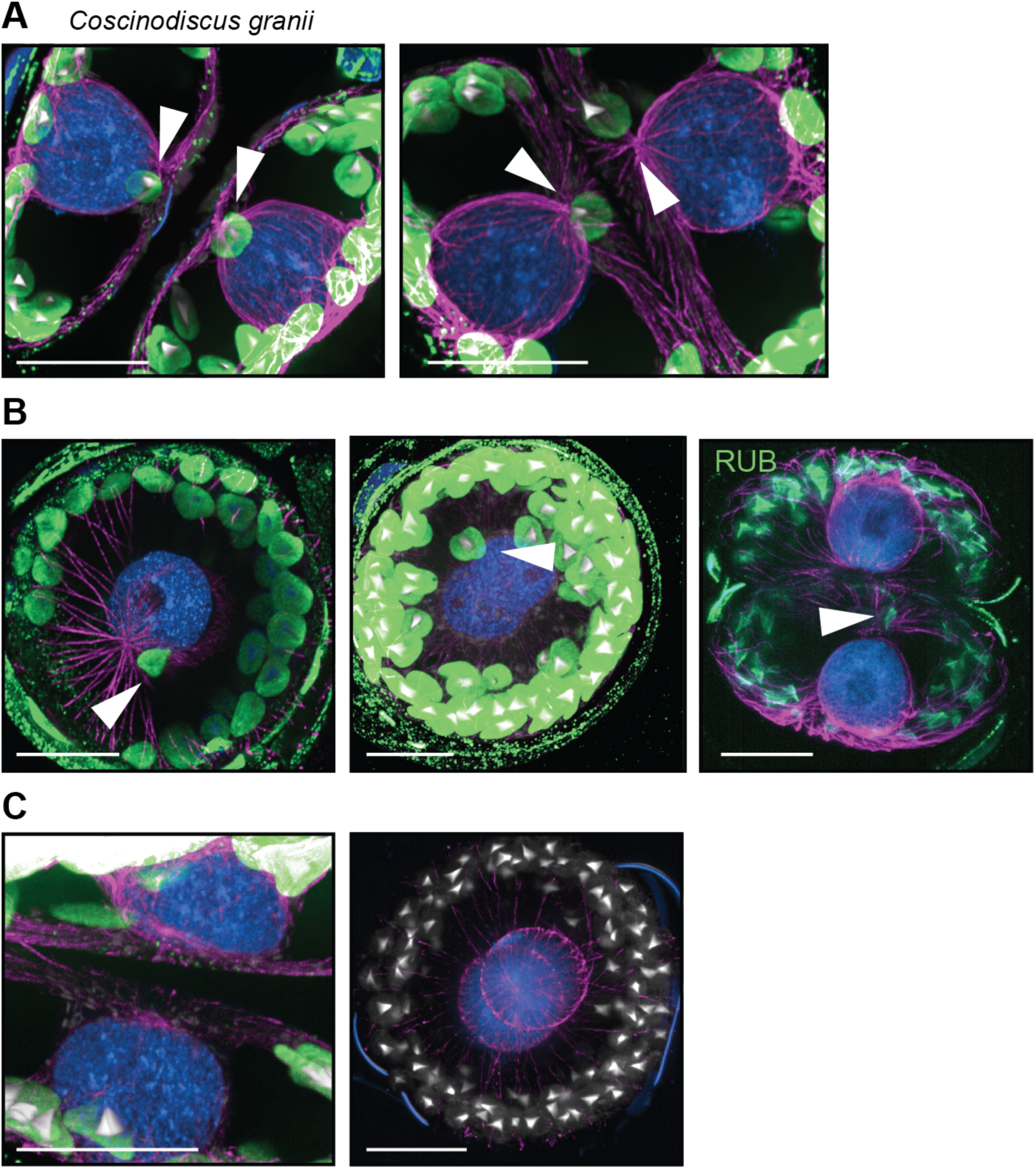
Chloroplast distribution in *Coscinodiscus granii* doublets. (A) Cells with single chloroplast localising close to the hypotheca in vicinity to the MTOC in both daughter cells. (B) Only one cell showing a hypothecal chloroplast, while (C) no chloroplast could be observed in MTOC proximity. Where not indicated differently, magenta indicates microtubules, blue DNA staining, grey NHS ester, and green PSII (rubisco staining in B panel 3). Scale bars represent 50 µm, biological scale 11.6 µm.

**Supplementary Figure 4:**
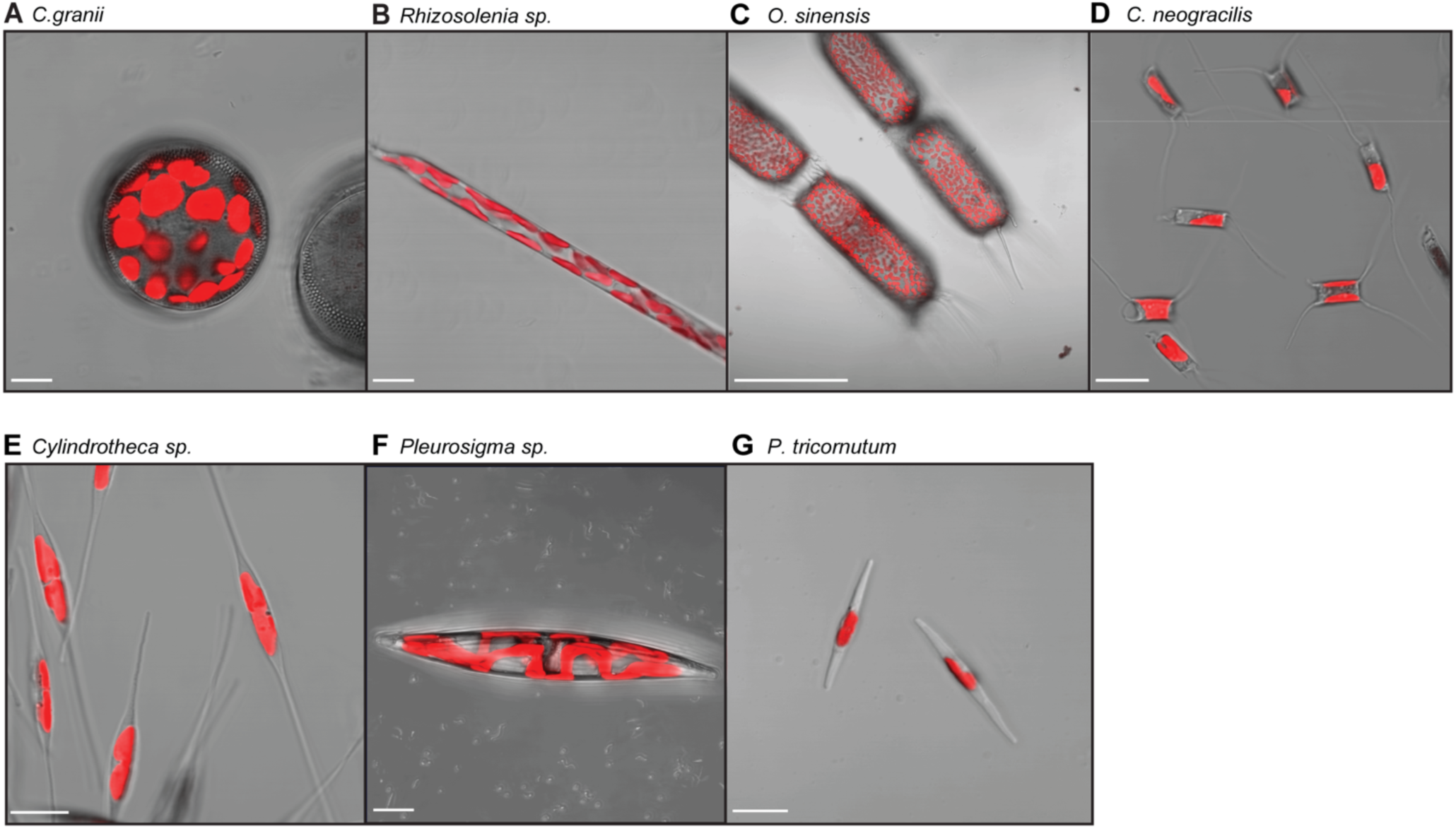
Confocal images of live cell cultures. Images using confocal ZEISS LSM 900 of (A) *C. granii* Scale 10 µm, (B) *Rhizosolenia sp.* Scale 10 µm, (C) *O. sinensis,* Scale 100 µm, (D) *C. neogracilis,* Scale 10 µm, (E) *Cylindrotheca sp*. Scale 10 µm, (F) *Pleurosigma sp*. max projection of the z-stack, Scale 10 µm, (G) *P. tricornutum,* Scale 10 µm.

**Supplementary Figure 5:**
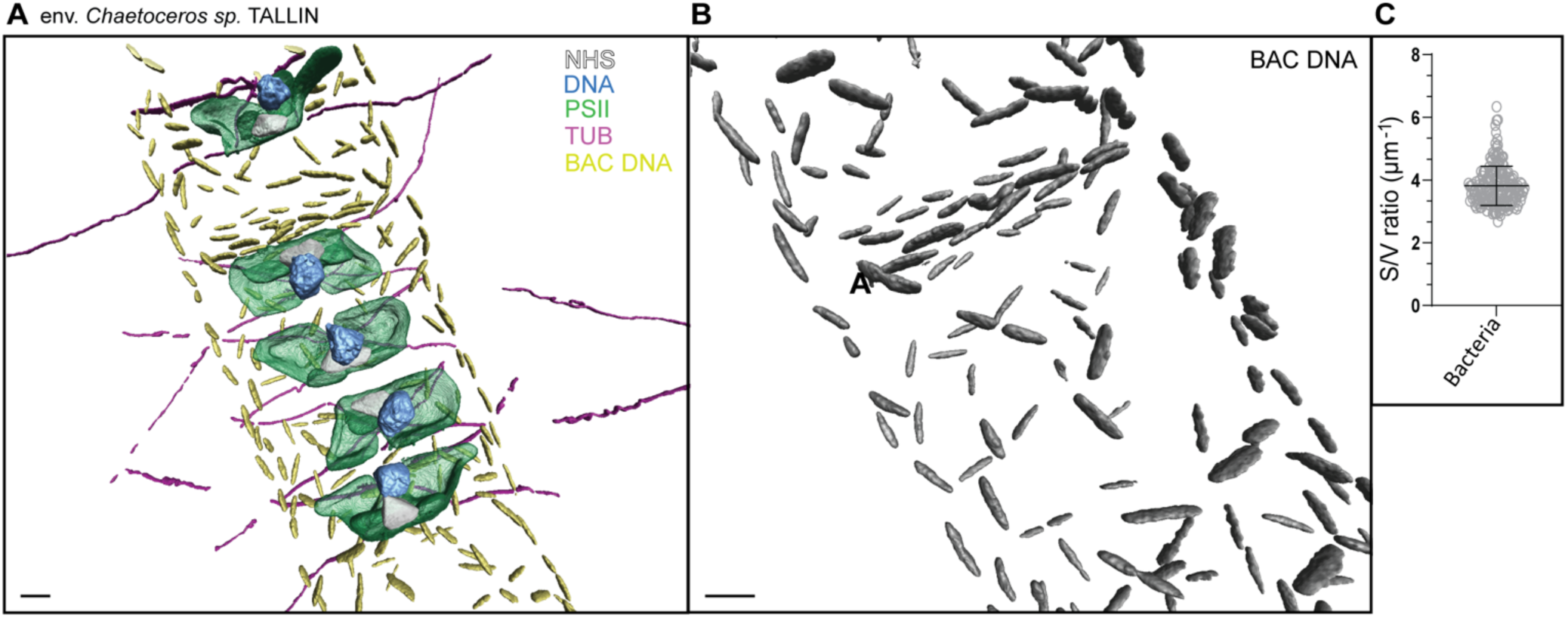
*Chaetoceros sp.* chain isolated in Tallinn, Estonia, with its associated microbiome. (A) 3D segmentation, expanded scale bar 10 µm, biological scale bar 2,5 µm. (B) Zoom on the segmented bacteria using DAPI staining expanded scale bar 10 µm biological scale bar 2,5 µm. (C) Surface to volume ratio of the bacteria (3.81 ± 0.63; n=186).

**Supplementary Figure 6:**
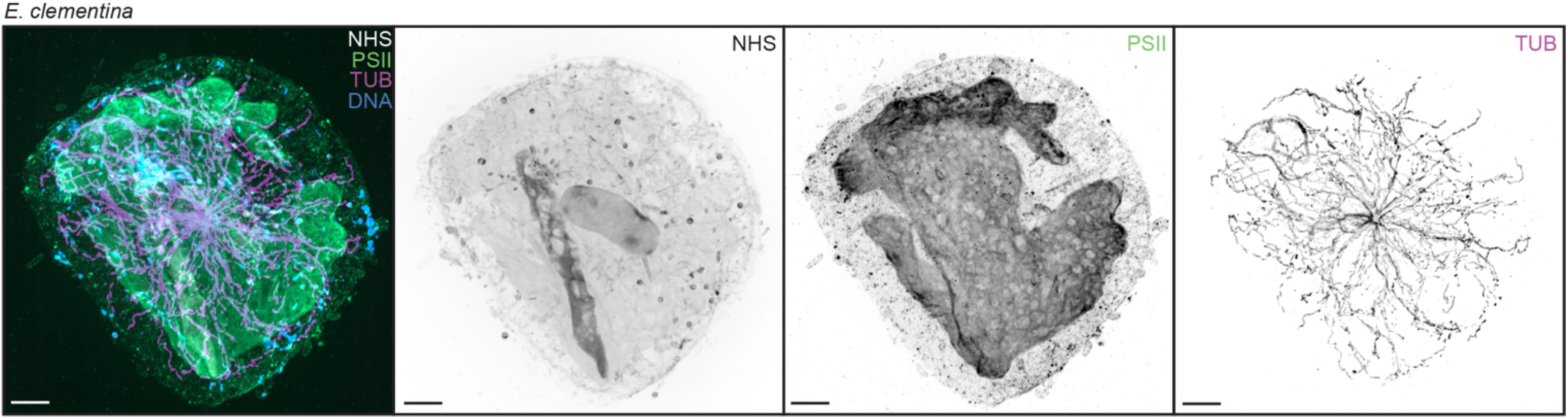
Interphase microtubules of *Epithemia clementina*. From left to right: a composite image showing NHS-Ester in grey, PSII in green, Tubulin in magenta, and DNA in blue. A greyscale image of just the NHS, PSII and Tubulin channels are on the right. Expanded scale bars are 10 µm and biological scale bars are 2.25 µm.

**Supplementary Figure 7:**
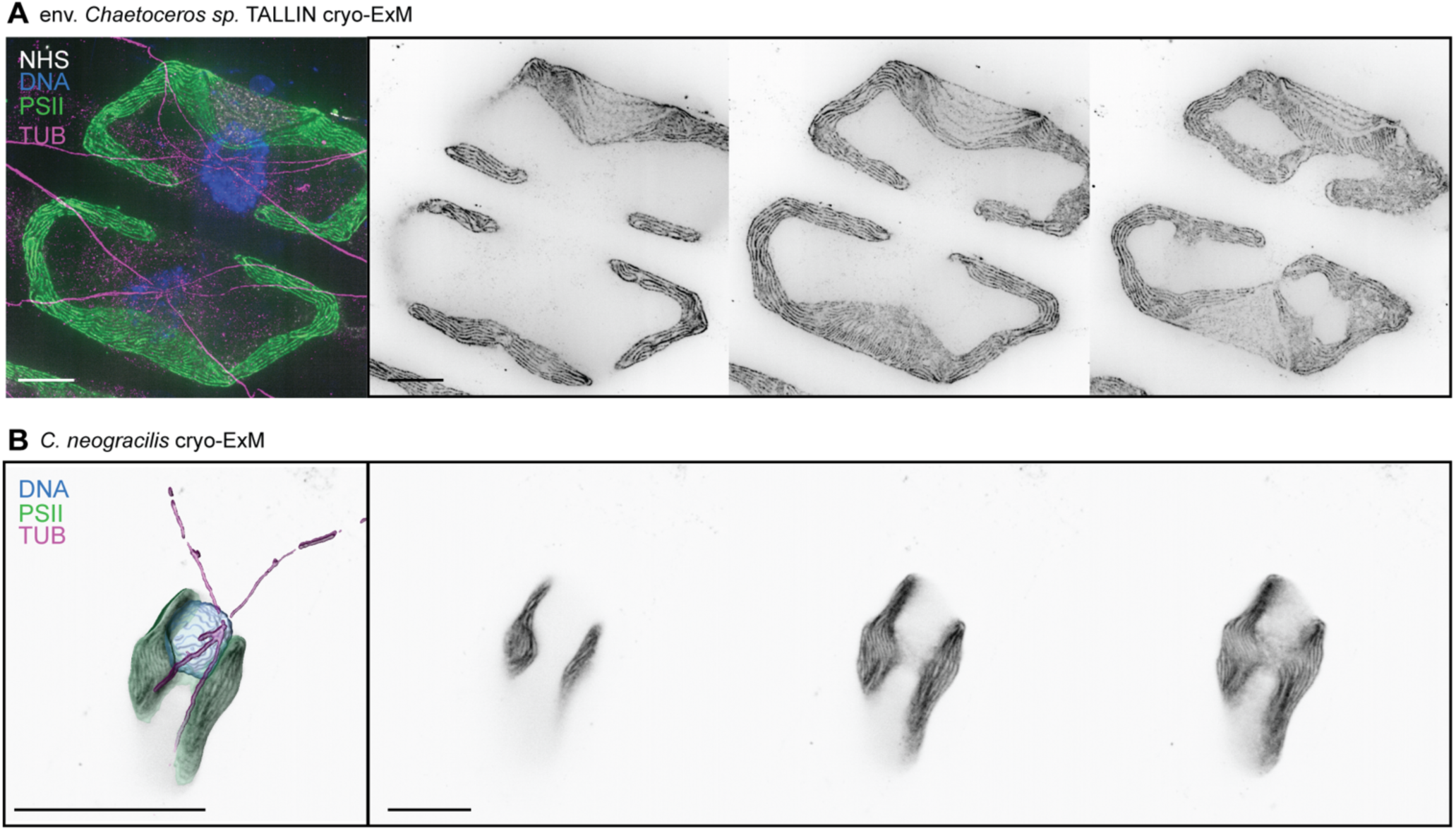
Chloroplasts of environmental *Chaetoceros sp.* and *C. neogracilis.* (A) Left panel is an overview of the env. *Chaetoceros sp.* isolated in Tallinn, Estonia, expanded scale bar 10 µm; biological scale bar 2.5 µm. PSII and NHS ester are shown as single slices, DNA and Tubulin are max projections. Three panels on the right are a montage of the z-stack showing the thylakoid membranes and PPTs. expanded scale bar 10 µm; biological scale bar 2.5 µm. (B) Left panel is a 3D segmentation of *C. neogracilis,* expanded scale bar 20 µm, biological scale bar 4.9 µm. Three panels on the right are a montage of the z-stack showing the thylakoid membranes and PPTs, expanded scale bar 10 µm, biological scale bar 2,46 µm.

## Supplementary table

**Supp. Table 1:**
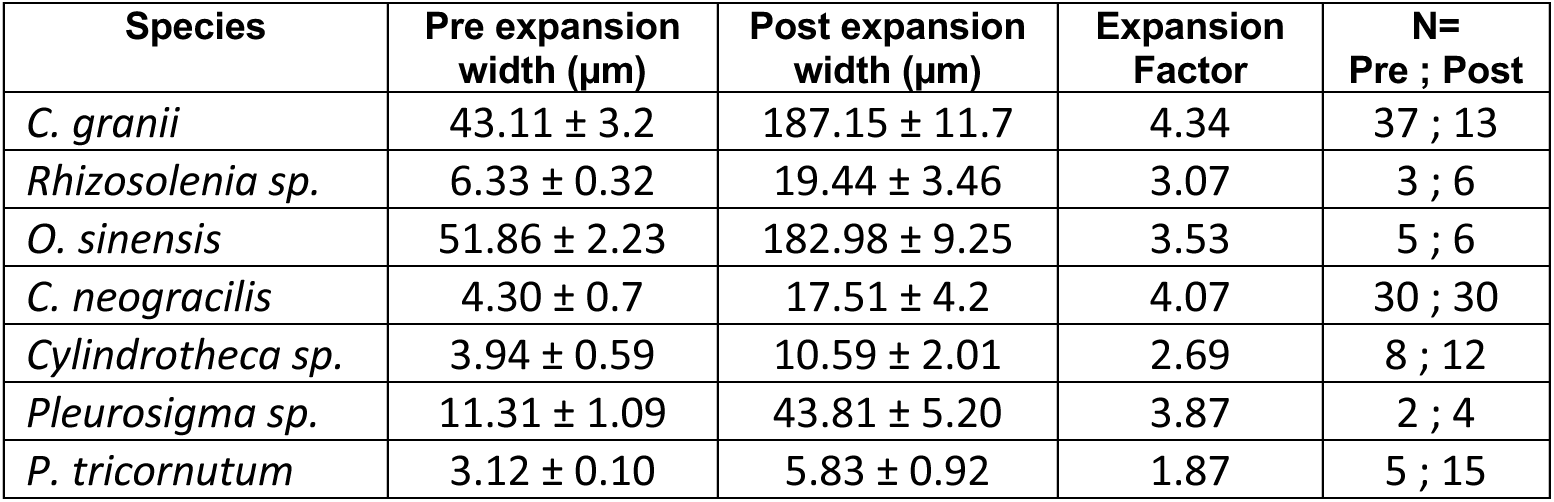
Species specific expansion factor in seven different diatom species. The expansion factor was calculated as described in (Gambarotto et al., 2019). For *C. neogracilis* and *C. granii*, pre-expanded measurements were done on gel. For all other species, direct measurements of the cells pre-expanded was done on fresh cultures. For all species, post-expanded measurements were performed in gel.

## Supplementary videos

**Supp. Movie 1:** Full volume of an expanded *Coscinodiscus granii* cell (**Fig. 1Di**) with the pyrenoid, labelled with NHS ester, shown in grey, PSII in green, microtubules in magenta and the nucleus in blue. The fragmented frustule is indicated in yellow, using DAPI signal for segmentation.

**Supp. Movie 2:** Full volume of an expanded environmental *Chaetoceros sp.* from Tallinn, Estonia, (**Fig. 1I**) with the pyrenoid, labelled with NHS ester, shown in grey, PSII in green, microtubules in magenta and the nucleus in blue. Bacteria colonising the surface are shown in cyan, using DAPI signal for segmentation.

**Supp. Movie 3:** Full volume of an expanded environmental *Chaetoceros sp.* chain from Plentzia, Spain (**Fig. 2E**, **Fig. 3B).** Using NHS ester, also labelling nuclear structures, the pyrenoids were segmented and are shown in grey, chloroplasts in green, microtubules in magenta and the nucleus in blue.

**Supp. Movie 4:** Close up of an expanded environmental *Chaetoceros sp.* chain from Plentzia, Spain (**Supp. Movie 3, Fig. 2E**, **Fig. 3B**). Thylakoid membranes are visible with PSII staining (green), with an individual membrane penetrating the pyrenoid (grey), labelled using NHS ester. Microtubule staining (magenta) reveals an extensive network and hollow MTOC.

**Supp. Movie 5:** Full volume of an expanded *Chaetoceros neogracilis* chain with the pyrenoid, labelled with NHS ester, shown in grey, PSII in green, microtubules in magenta and the nucleus in blue.

